# Genome-wide identification of a regulatory mutation in *BMP15* controlling prolificacy in sheep

**DOI:** 10.1101/858860

**Authors:** Louise Chantepie, Loys Bodin, Julien Sarry, Florent Woloszyn, Florence Plisson-Petit, Julien Ruesche, Laurence Drouilhet, Stéphane Fabre

**Author notes:** corresponding author (SF).

## Abstract

The search for the genetic determinism of prolificacy variability in sheep has evidenced several major mutations in genes playing a crucial role in the control of ovulation rate. In the Noire du Velay (NV) sheep population, a recent genetic study has evidenced the segregation of such a mutation named *FecL*^*L*^. However, based on litter size (LS) records of *FecL*^*L*^ non-carrier ewes, the segregation of a second prolificacy major mutation was suspected in this population. In order to identify this mutation, we have combined case/control genome-wide association study with ovine 50k SNP chip genotyping, whole genome sequencing and functional analyses. A new single nucleotide polymorphism (OARX:50977717T>A, NC_019484) located on the X chromosome upstream of the *BMP15* gene was evidenced highly associated with the prolificacy variability (*P* =1.93E^-11^). The variant allele was called *FecX*^*N*^ and shown to segregate also in the Blanche du Massif Central (BMC) sheep population. In both NV and BMC, the *FecX*^*N*^ allele frequency was estimated close to 0.10, and its effect on LS was estimated at +0.20 lamb per lambing at heterozygous state. Homozygous *FecX*^*N*^ carrier ewes were fertile with increased prolificacy in contrast to numerous mutations affecting *BMP15*. At the molecular level, *FecX*^*N*^ was shown to decrease BMP15 promoter activity and to impact *BMP15* expression in oocyte. This regulatory action was proposed as the causal mechanism for the *FecX*^*N*^ mutation to control ovulation rate and prolificacy in sheep.

**Author Summary:** In the genetic etiology of women infertility syndromes, a focus was done on the oocyte-expressed *BMP15* and *GDF9* genes harboring several mutations associated with ovarian dysfunctions. In sheep also, mutations in these two genes are known to affect the ovarian function leading to sterility or, on the opposite, increasing ovulation rate and litter size constituting the prolificacy trait genetically selected in this species. Through a genome-wide association study with the prolificacy phenotype conducted in the French Noire du Velay sheep breed, we describe a novel mutation located in the regulatory region upstream of the *BMP15* gene on the X chromosome. This mutation increases litter size by +0.2 lamb per lambing at the heterozygous state, possibly through an inhibition of *BMP15* expression within the oocyte. Our findings suggest a novel kind of *BMP15* variant responsible for high prolificacy, in contrast to all other *BMP15* variants described so far in the coding sequence.

## Introduction

There is now an accumulation of evidence that oocyte plays a central role in controlling the ovarian folliculogenesis, from the early stages up to ovulation. Among the local factors produced by the oocyte itself, members of the bone morphogenetic protein/growth and differentiation factor (BMP/GDF) family play an integral role in this control (Persani et al., 2015[1]). Among them, the most important are surely *BMP15* and *GDF9*. Knock-out mice models gave the first evidence of the importance of these two oocyte-derived factors acting individually as homodimers and/or through a synergistic co-operation to control the ovarian function (Elvin et al., 1999, Yan et al., 2001[2,3]).

In human also, a focus was done on *BMP15* and *GDF9* about their implication in various ovarian dysfunctions. Indeed, numerous heterozygous missense mutations have been identified in both genes associated with primary or secondary amenorrhea in different cohorts of women affected by primary ovarian insufficiency (POI) all over the world. Particularly, the 10-fold higher prevalence of *BMP15* variants among patients with POI compared with the control population supports the causative role of these mutations (Persani et al., 2015[1]). Alteration of *BMP15* and *GDF9* were also searched in association with the polycystic ovary syndrome (PCOS). Here again several missense variants were discovered in both genes, but the pathogenic role of these mutations remains controversial in the etiology of this syndrome. However, several studies have reported an aberrant expression of *BMP15* and *GDF9* in the ovary of PCOS patients (Teixera Filho et al. 2002; Wei et al. 2014 [4,5]). Interestingly, some *BMP15* polymorphisms situated in the 5’UTR are significantly associated with the over response to recombinant FSH applied during assisted reproductive treatment and with the risk to develop an ovarian hyperstimulation syndrome (OHSS, Moron et al. 2006; Hanevik et al. 2011 [6,7]). Finally, polymorphisms in *BMP15* and *GDF9* genes were also searched in association with dizygotic twinning in human. If no convincing results were obtained for *BMP15*, some lost-of-function variants of *GDF9* were observed significantly more frequently in mothers of twins compared to the control population (Palmer et al. 2006; Simpson et al. 2014 [8,9]).

In parallel, the search for the genetic determinism of ovulation rate and prolificacy variability in sheep has also highlighted the crucial role of *BMP15* and *GDF9* by evidencing numerous independent loss-of-function mutations all altering the coding sequence of these two genes (Persani et al. 2015; Abdoli et al. 2016 [1,10]). Depending on the mutation and its hetero- or homozygous state, the phenotype controlled by these mutations in *BMP15* and *GDF9* goes from the early blockade of the folliculogenesis, and subsequent sterility, to an extraordinary increase of the ovulation rate (OR) and thus litter size (LS) of carrier ewes (Galloway et al., 2000; Hanrahan et al., 2004; Sylva et al., 2011; Demars et al., 2013 [11–14]). Thus, sheep exhibiting an extremely high prolificacy are of great interest for identifying genes and mutations involved in molecular pathways controlling the ovarian function. These animal models have a double interest, in agriculture for the genetic improvement of the prolificacy, and in human clinic for providing valuable candidate genes in the genetic determinism of female infertility or subfertility, as described above.

The Noire du Velay (NV) population is a French local sheep breed mainly reared in the Haute-Loire and Loire departments. Ewes present naturally out-of-season breeding ability, very good maternal characteristics and a quite high prolificacy (mean LS=1.62 lamb per lambing). Large variation in LS has been observed in this breed and a recent genetic study has evidenced the segregation of an autosomal mutation named *FecL*^*L*^ controlling this trait (Chantepie et al. 2018[15]). This variant located in the intron 7 of the *B4GALNT2* gene and associated with its ectopic ovarian expression, was originally discovered in the Lacaune meat sheep breed, increasing OR and prolificacy (Drouilhet et al. 2013[16]). For the segregation study, more than 2700 NV ewes with LS records were genotyped at the *FecL* locus (Chantepie et al. 2018[15]). Surprisingly, the distribution of LS and the existence of high prolific ewes among the *FecL*^*L*^ non-carrriers have suggested the possible segregation of a second prolificacy major mutation in this population as already observed in the Lacaune breed carrying both *FecL*^*L*^ and *FecX*^*L*^ (Bodin et al. 2007, Drouilhet et al. 2013[16,17]). In order to validate this hypothesis, after specific genotyping excluding all other known mutations affecting OR and LS and segregating in French sheep populations, we have performed a genome-wide association study (GWAS) based on a case/control design. Completed by the whole genome sequencing of two finely chosen animals, we have identified a new regulatory variant called *FecX*^*N*^ affecting the oocyte-dependent expression of *BMP15* in association with increased prolificacy in sheep.

## Results

### Genetic association analyses

A first set of genomic DNA from 30 NV ewes without the *FecL*^*L*^ prolific allele at the *B4GALNT2* locus (LS records ranging from 2.00 to 3.00) was genotyped for already known mutations affecting sheep prolificacy at the 3 other loci, *BMPR1B, GDF9* and *BMP15*. Using specific RFLP assay (*BMPR1B*, Wilson et al. 2001[18]) or Sanger sequencing of coding parts (*GDF9* and *BMP15*, Talebi et al, 2018[19]), none of the known mutations were evidenced (data not shown). Thus, to establish the genetic determinism of the remaining LS variation in this population, 80 ewes were genotyped by Illumina Ovine SNP50 Genotyping Beadchip. The allele frequencies of the most highly prolific ewes (cases, n=40, mean LS=2.47) and lowly prolific ewes (controls, n=40, mean LS=1.23) were compared to identify loci associated with LS using GWAS according to the procedures described in the Materials and Methods. Finally, genotype data were obtained from 79 animals (39 cases, 40 controls). Six markers located on OARX were significantly associated with LS variation at the genome-wide level after Bonferroni correction (Fig 1A, Table 1). Importantly, at the chromosome-wide level, a cluster of 26 significant markers encompassed the location of the *BMP15* candidate gene (Fig 1B). In order to better characterize this locus on the X chromosome, we have determined for each individual the most likely linkage phase across 80 markers (10Mb) including the significant region. After haplotype clusterization, a specific segment of 3.5 Mb (50639087–54114793 bp, OARv3.1 genome assembly) was identified to be more frequent in highly prolific cases than in controls (f_cases_= 0.51 vs. f_controls_=0.37, *P*= 1.92E^-11^, Chi-square test) (Fig 2). This identified segment contained the *BMP15* gene (50970938-50977454 bp, OARv3.1) well-known to play a crucial role in the ovarian function and to be a target of numerous mutations in its coding region controlling prolificacy (Persani et al, 2015[1]).

**Table 1.** Markers significantly associated with litter size.

| SNP | Chromosome | Position<sup>a</sup> | MAF<sup>b</sup> | $P_{\text{Unadj}}$<sup>c</sup> | $P_{\text{Chrom}}$<sup>d</sup> | $P_{\text{Genome}}$<sup>e</sup> |
| --- | --- | --- | --- | --- | --- | --- |
| OARX_51294776.1 | OARX | 53756339 | 0.28 | 3.52E<sup>-11</sup> | 4.18E<sup>-08</sup> | 1.67E<sup>-06</sup> |
| s27837.1 | OARX | 53825247 | 0.30 | 5.83E<sup>-10</sup> | 6.92E<sup>-07</sup> | 2.77E<sup>-05</sup> |
| s73460.1 | OARX | 53905939 | 0.44 | 9.59E<sup>-10</sup> | 1.14E<sup>-06</sup> | 4.55E<sup>-05</sup> |
| s39212.1 | OARX | 53852735 | 0.44 | 3.46E<sup>-09</sup> | 4.11E<sup>-06</sup> | 1.64E<sup>-04</sup> |
| OARX_52608221.1 | OARX | 52367253 | 0.41 | 1.68E<sup>-08</sup> | 2.00E<sup>-05</sup> | 7.99E<sup>-04</sup> |
| s46003.1 | OARX | 80222479 | 0.44 | 2.56E<sup>-07</sup> | 3.03E<sup>-04</sup> | 1.21E<sup>-02</sup> |
| OARX_55032299.1 | OARX | 48942926 | 0.30 | 1.57E<sup>-06</sup> | 1.86E<sup>-03</sup> | NS |
| OARX_72164491.1 | OARX | 74388397 | 0.23 | 1.71E<sup>-06</sup> | 2.03E<sup>-03</sup> | NS |
| OARX_49135019.1 | OARX | 42475099 | 0.35 | 1.80E<sup>-06</sup> | 2.13E<sup>-03</sup> | NS |
| OARX_111306030.1 | OARX | 92041520 | 0.27 | 2.25E<sup>-06</sup> | 2.67E<sup>-03</sup> | NS |
| OARX_102620828.1 | OARX | 82796975 | 0.41 | 3.02E<sup>-06</sup> | 3.59E<sup>-03</sup> | NS |
| s31917.1 | OARX | 58202482 | 0.44 | 3.12E<sup>-06</sup> | 3.70E<sup>-03</sup> | NS |
| OARX_72351736.1 | OARX | 74590448 | 0.19 | 5.63E<sup>-06</sup> | 6.68E<sup>-03</sup> | NS |
| OARX_72263548.1 | OARX | 74498463 | 0.17 | 6.45E<sup>-06</sup> | 7.66E<sup>-03</sup> | NS |
| OARX_49564109.1 | OARX | 42876169 | 0.49 | 8.16E<sup>-06</sup> | 9.69E<sup>-03</sup> | NS |
| DU400878_520.1 | OARX | 73847207 | 0.27 | 9.96E<sup>-06</sup> | 1.18E<sup>-02</sup> | NS |
| s54281.1 | OARX | 58993959 | 0.24 | 1.20E<sup>-05</sup> | 1.42E<sup>-02</sup> | NS |
| s05229.1 | OARX | 58346644 | 0.46 | 1.71E<sup>-05</sup> | 2.03E<sup>-02</sup> | NS |
| s27938.1 | OARX | 53275559 | 0.35 | 1.77E<sup>-05</sup> | 2.10E<sup>-02</sup> | NS |
| OARX_54104393.1 | OARX | 49870983 | 0.36 | 2.03E<sup>-05</sup> | 2.41E<sup>-02</sup> | NS |
| OARX_72236232.1 | OARX | 74464263 | 0.18 | 2.23E<sup>-05</sup> | 2.64E<sup>-02</sup> | NS |
| OARX_111349974.1 | OARX | 92085118 | 0.18 | 2.23E<sup>-05</sup> | 2.65E<sup>-02</sup> | NS |
| OARX_53703822.1 | OARX | 51193144 | 0.35 | 2.59E<sup>-05</sup> | 3.08E<sup>-02</sup> | NS |
| OARX_43227227.1 | OARX | 36235514 | 0.32 | 3.41E<sup>-05</sup> | 4.04E<sup>-02</sup> | NS |
| OARX_51842287.1 | OARX | 53162079 | 0.49 | 3.43E<sup>-05</sup> | 4.08E<sup>-02</sup> | NS |
| OARX_102654502.1 | OARX | 82837158 | 0.20 | 3.65E<sup>-05</sup> | 4.33E<sup>-02</sup> | NS |

<sup>a</sup> Position of markers are based on the OARv3.1 assembly in bp.
<sup>b</sup> MAF, minor allele frequency.
<sup>c</sup> $P_{\text{Unadj}}$ corresponds to exact unadjusted p-value for the Fisher's test.
<sup>d</sup> $P_{\text{Chrom}}$ corresponds to p-value after chromosome-wide Bonferroni correction.
<sup>e</sup> $P_{\text{Genome}}$ corresponds to p-value after genome-wide Bonferroni correction (NS, non-significant).

**Figure 1.**
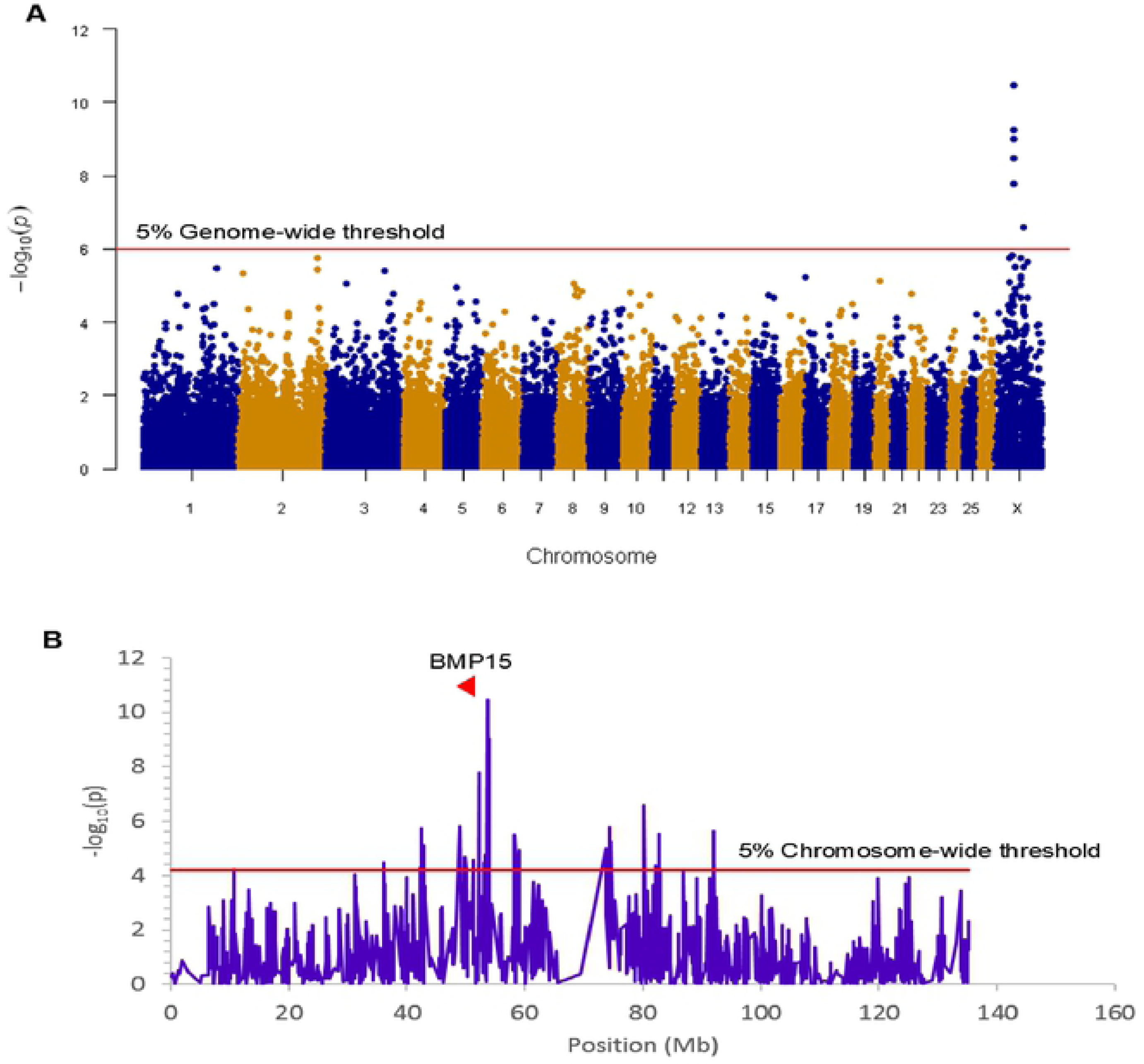
Genome-wide and chromosome-wide association results. (A) Genome-wide association results for litter size in the NV sheep population. Manhattan plot shows the combined association signals (-log_10_(p-value)) on the y-axis versus SNPs position in the sheep genome on the x-axis and ordered by chromosome number (assembly OARv3.1). Red line represents the 5% genome-wide threshold. (B) OARX chromosome-wide association results. The curve shows the combined association signals (-log_10_(p-value)) on the y-axis versus SNPs position on the X chromosome on the x-axis (assembly OARv3.1). Red line represents the 5% chromosome wide threshold. The *BMP15* gene location is indicated by a red arrowhead.

**Figure 2.**
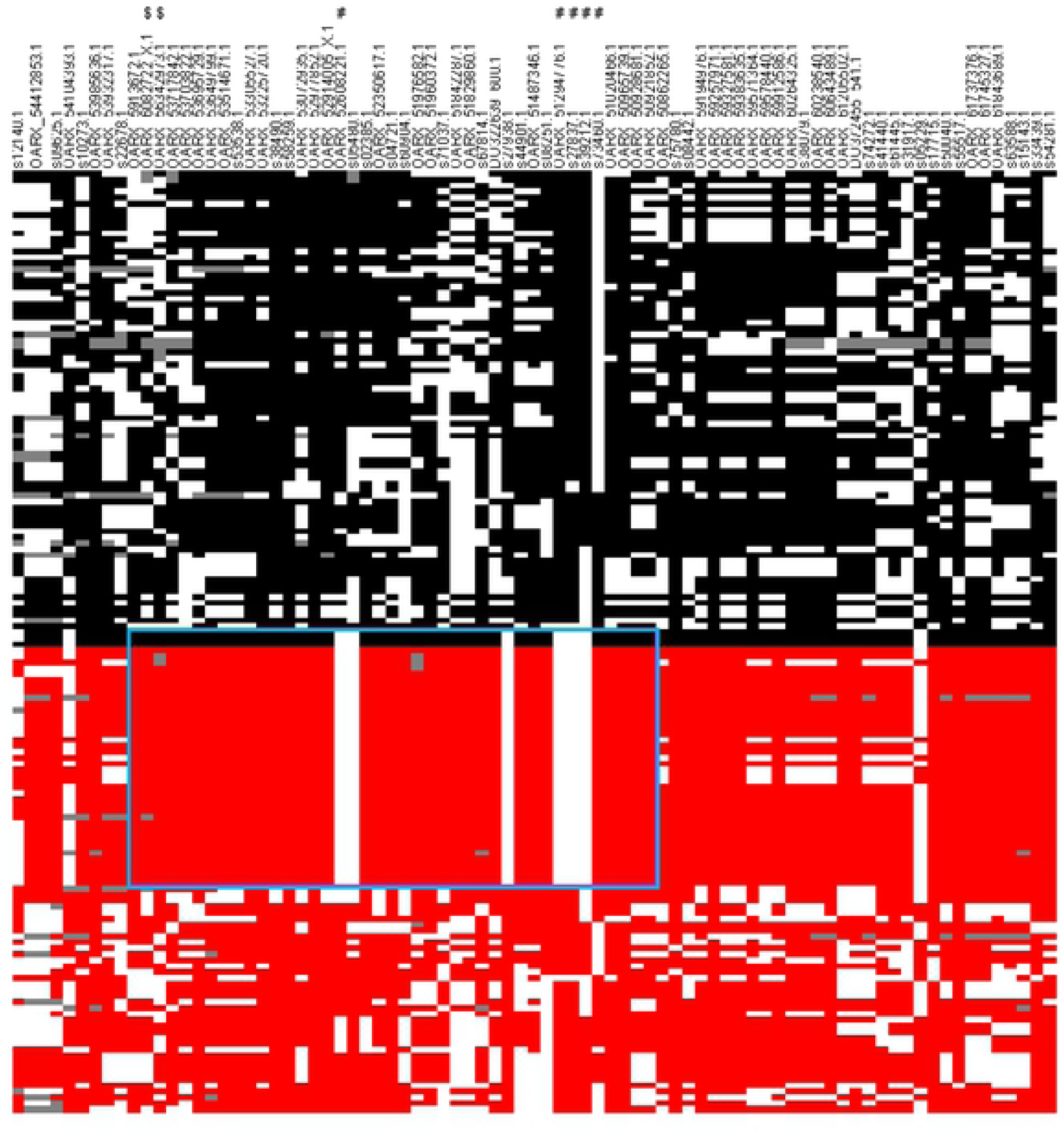
Clusterization of haplotypes reconstructed at the OARX locus. 80 markers encompassing the OARX region of interest (50.6 Mb-54.1 Mb) were selected to construct haplotypes from 39 cases and 40 control animals. Each column represents one SNP and each line represents one haplotype. For one marker (i) allele 1 is in black in controls, or in red in cases, (ii) allele 2 is in white when the phase was unambiguous and (iii) grey color represents unphased SNP. Haplotypes were ordered to distinguish controls versus cases and clusterized to classify similar clades of haplotypes. The # sign flags SNP significantly associated with LS at genome-wide level. The $ sign flags SNP flanking the *BMP15* genes (50970938-50977454 bp). The specific haplotype preferentially selected in highly prolific ewes (cases) is symbolized by the cyan rectangle.

### Characterization of the mutation

While the *BMP15* gene could be considered as a positional and functional candidate gene, no mutation was evidenced by Sanger sequencing of the *BMP15* coding regions of the most prolific ewes studied. In order to find the potential causal mutation, we sequenced the whole genome of two finely chosen ewes based on the shortest haplotype within the region (homozygous reference vs. homozygous variant) and their opposite extreme phenotypes (LS 1.1 vs. 2.8).

Variant search analysis and annotations through GATK toolkit was limited to the OARX: 50639087-54114793 region. We detected 60 SNPs and 90 small insertions and deletions (INDELs) with quality score >30 (S1 Table). Among them, we particularly focused on the 85 variants located within annotated genes (upstream, exon, intron, splice acceptor or donor, and downstream localization). After filtering these 85 variants for allele sharing with other breeds based on SheepGenome DB (http://sheepgenomesdb.org/) and 68 publicly available domestic sheep genomes (International Sheep Genomics Consortium; http://www.sheephapmap.org/), none of them were removed, all being NV breed specific. Finally, and based on prolificacy gene knowledge, we were particularly interested in one SNP (T>A) identified in the upstream region of the *BMP15* gene at position 50977717 on OARX v3.1. We then developed a RFLP assay to specifically genotype for this polymorphism. Among the 79 animals of the GWAS, 31 ewes were heterozygous and 6 homozygous for the A variant allele. As shown in Table 2, most of the A carrier ewes were in the highly prolific Case group (34 among 39), while only 3 set in the Control group. When associating the LS performance of the 79 ewes to their genotype at the OARX: 50977717T>A SNP, the A non-carriers exhibited a mean LS of 1.36, heterozygous T/A a mean LS of 2.32 and homozygous A/A a mean LS of 2.73 indicating that the A allele of this polymorphism was strongly associated with increased LS in NV (T/A or A/A vs. T/T, *P*<1E^-3^, one-way ANOVA). Furthermore, this polymorphism appears in total linkage disequilibrium with the six more significant markers from the GWAS analysis (Fig 3). Genotype information at the OARX: 50977717T>A locus was introduced in the GWAS analysis. This SNP appeared as the most significant marker associated to the prolificacy phenotype (*P*_unadjusted_=1.93E^-11^, *P*_Chromosome-wide corrected_=1.62E^-14^ and *P*_Genome-wide corrected_ =9.13E^-07^) suggesting that it could be the causal mutation (S3 Figure). In accordance with the *Fec* gene nomenclature, the mutant allele identified upstream of the *BMP15* gene in NV sheep was named *FecX*^*N*^.

**Table 2.** Distribution of OARX:50977717T>A SNP genotypes and associated LS in case and control groups.

| Group |  | TT | TA | AA |
| --- | --- | --- | --- | --- |
| Low prolific control | n= | 37 | 3 |  |
|  | Raw mean LS | 1.22 | 1.39 |  |
| Highly prolific case | n= | 5 | 28 | 6 |
|  | Raw mean LS | 2.41 | 2.43 | 2.73 |
| **Total** |  | **42** | **31** | **6** |

**Figure 3.**
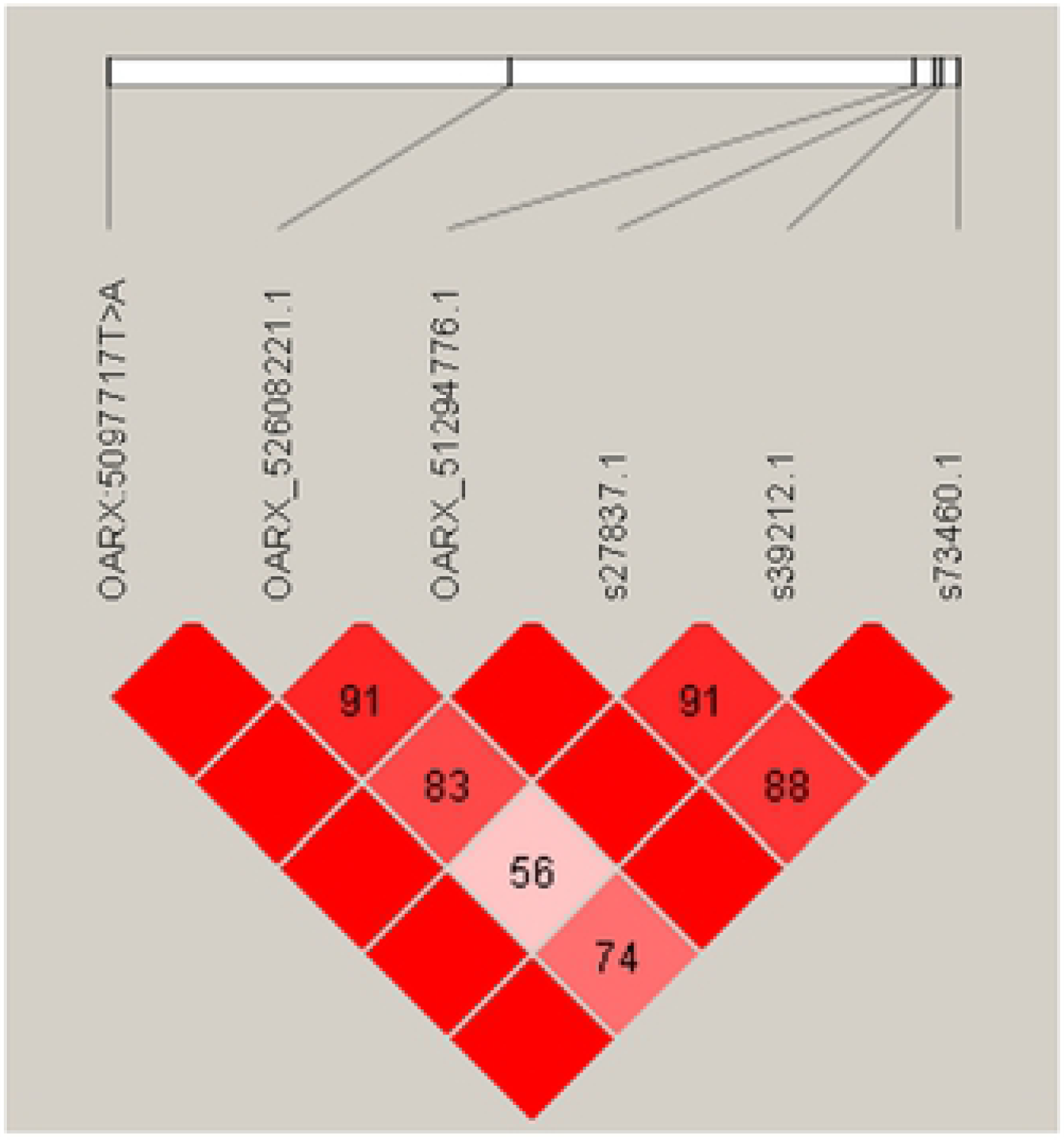
Linkage disequilibrium (LD) plot. Generated by Haploview, pair-wise LD between SNP markers (OARX: 50977717T>A and the five genome-wide significant LS-associated markers) is represented by the D’ value. Strong LD is represented in dark red boxes (D’=100) and weaker LD (D’<100) in lighter red boxes.

As described for other prolific alleles such as *FecB*^*B*^, *FecX*^*G*^, *FecG*^*H*^, *FecX*^*Gr*^ and *FecL*^*L*^, a given mutation can segregate in several sheep populations (Davis et al., 2002; Mullen et al., 2013, Chantepie et al. 2018, Ben Jemaa et al. 2019[15,20–22]). We have tested the *FecX*^*N*^ allele presence in a diversity of 26 sheep breeds representing 725 animals (Rochus et al. 2018[23]). Among the breeds tested, the *FecX*^*N*^ genotyping has confirmed the segregation of this mutation in NV breed and revealed its presence in the Blanche du Massif Central (BMC) and Lacaune breeds (Table 3). Additionally, the *FecX*^*N*^ variant was absent from the Ensembl variant database (http://www.ensembl.org) compiling information from i) dbSNP, ii) whole genome sequencing information from the NextGen project (180 animals from various Iranian and Moroccan breeds) and iii) the International Sheep Genome Consortium (551 animals from 39 breeds all over the world).

**Table 3.** *FecX*^*N*^ genotype distribution from a diversity panel of French ovine breeds.

| Breed | Total | Genotype |  | Breed | Total | Genotype |  |
| --- | --- | --- | --- | --- | --- | --- | --- |
|  |  | +/+ | N/+ |  |  | +/+ | N/+ |
|  |  | (+/Y) | (N/Y)<sup>a</sup> |  |  | (+/Y) | (N/Y)<sup>a</sup> |
| Berrichon du Cher | 29 | 29 |  | Mourerous | 26 | 26 |  |
| Blanche Massif Central | 31 | 27 | 4 | Mouton Vendéen | 30 | 30 |  |
| Causse du Lot | 32 | 32 |  | Noire du Velay | 28 | 26 | 2 |
| Charmoise | 31 | 31 |  | Préalpes du sud | 27 | 27 |  |
| Charollais | 29 | 29 |  | Rava | 29 | 29 |  |
| Corse | 30 | 30 |  | Romane | 29 | 29 |  |
| Ile de France | 28 | 28 |  | Romanov | 26 | 26 |  |
| Lacaune (meat) | 42 | 40 | 2 | Rouge de l'Ouest | 28 | 28 |  |
| Lacaune (dairy) | 40 | 40 |  | Roussin | 30 | 30 |  |
| Limousine | 30 | 30 |  | Suffolk | 20 | 20 |  |
| Manech tête rousse | 29 | 29 |  | Tarasconnaise | 32 | 32 |  |
| Martinik | 22 | 22 |  | Texel | 21 | 21 |  |
| Merinos d'Arles | 26 | 26 |  |  |  |  |  |
| TOTAL |  |  |  |  | 725 | 717 | 8 |

<sup>a</sup>: +/+ or N/+ females, N/Y or N/+ hemizygous males.

### *FecX*^*N*^ genotype frequency and effect on prolificacy

Large cohorts of ewes, chosen at random, were genotyped in order to accurately estimate the allele frequencies in the NV and the BMC populations (Table 4). The frequency of the N prolific allele at the *FecX* locus was similar in both populations, 0.11 and 0.10, with a distribution of 19.4% and 17.6% heterozygous, 1.5 % and 1% homozygous carriers in NV and BMC, respectively. The genotype frequencies were consistent with the Hardy Weinberg equilibrium (HWE) in both breed (NV *P*= 0.28 and BMC *P*= 0.76).

**Table 4.**
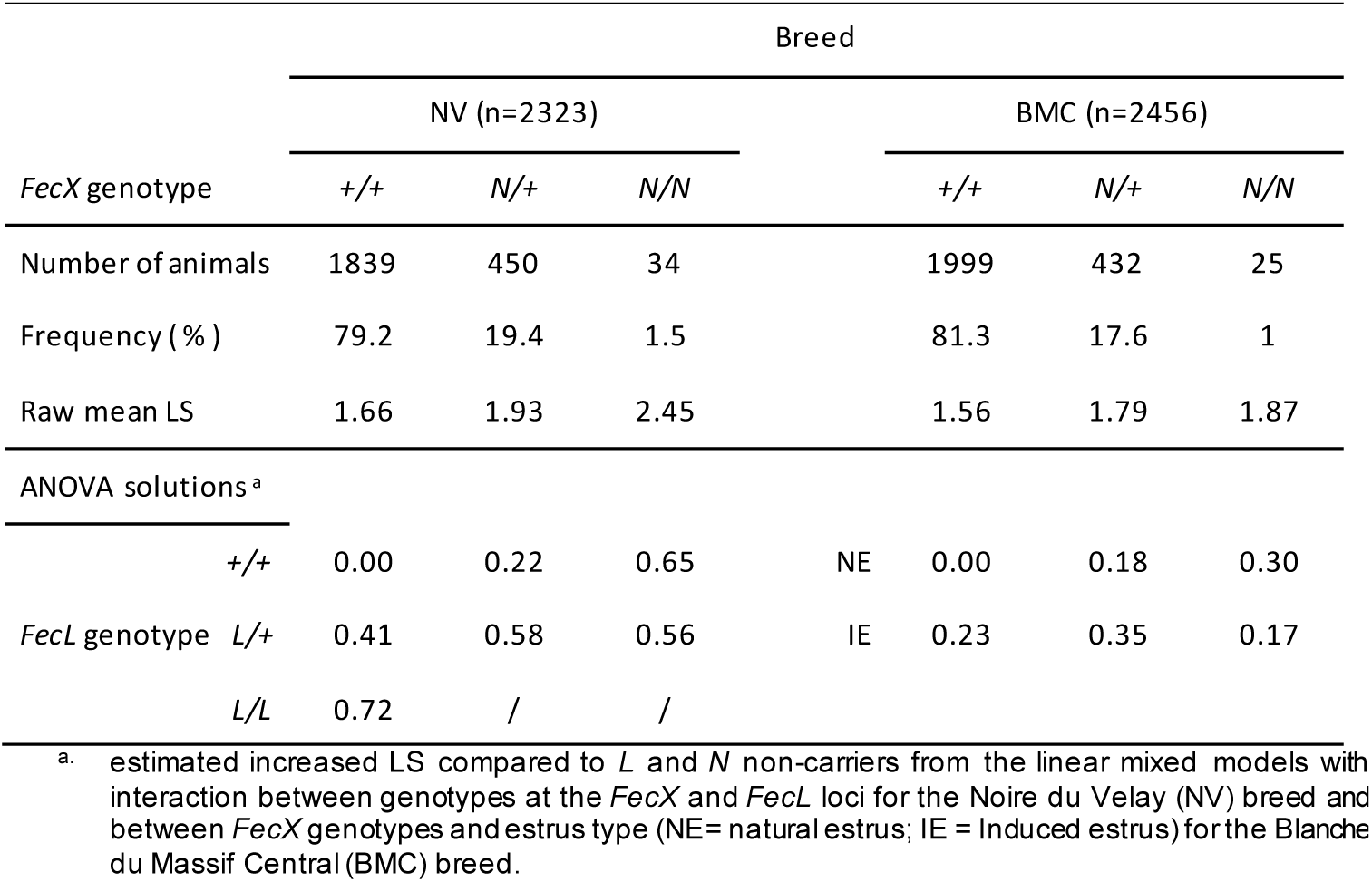
*FecX*^*N*^ frequencies and effects on LS in NV and BMC breeds.

Based on the raw mean LS observations, the *FecX*^*N*^ carrier ewes clearly exhibited increased LS compared to non-carriers in both populations (Table 4). The L prolific allele at the *FecL* locus is also segregating in NV (Chantepie et al. 2018 [15]). Results of the linear mixed model showed that for the NV breed, one copy of the *FecX*^*N*^ allele significantly increased LS by +0.22 and two copies increased LS by +0.65, while a single copy of the *FecL*^*L*^ allele increased LS by +0.41 and two copies by +0.72. Based on the 80 ewes genotyped heterozygous at both loci it appeared that the effect of *FecX*^*N*^ and *FecL*^*L*^ on LS was not fully additive, the expected LS being significantly slightly reduced by −0.05 (0.58 instead of 0.63) (Fig. 4A). For the BMC population, compared to *FecX*^*+*^*/FecX*^*+*^ ewes, *FecX*^*N*^*/FecX*^*+*^ exhibited increased LS by +0.18 and *FecX*^*N*^*/FecX*^*N*^ by +0.30 under natural estrus (Fig. 4B). The use of PMSG for estrus synchronization increased LS significantly among *FecX*^*+*^*/FecX*^*+*^ ewes (+0.23) and *FecX*^*N*^*/FecX*^*+*^ ewes (+0.18) while the effect on *FecX*^*N*^*/FecX*^*N*^ ewes was negative (−0.13). The combined effect of the first copy of the *FecX*^*N*^ allele and the use of PMSG treatment was not fully additive, the interaction being significant although low (0.35 instead of 0.41) (Fig. 4B).

**Figure 4.**
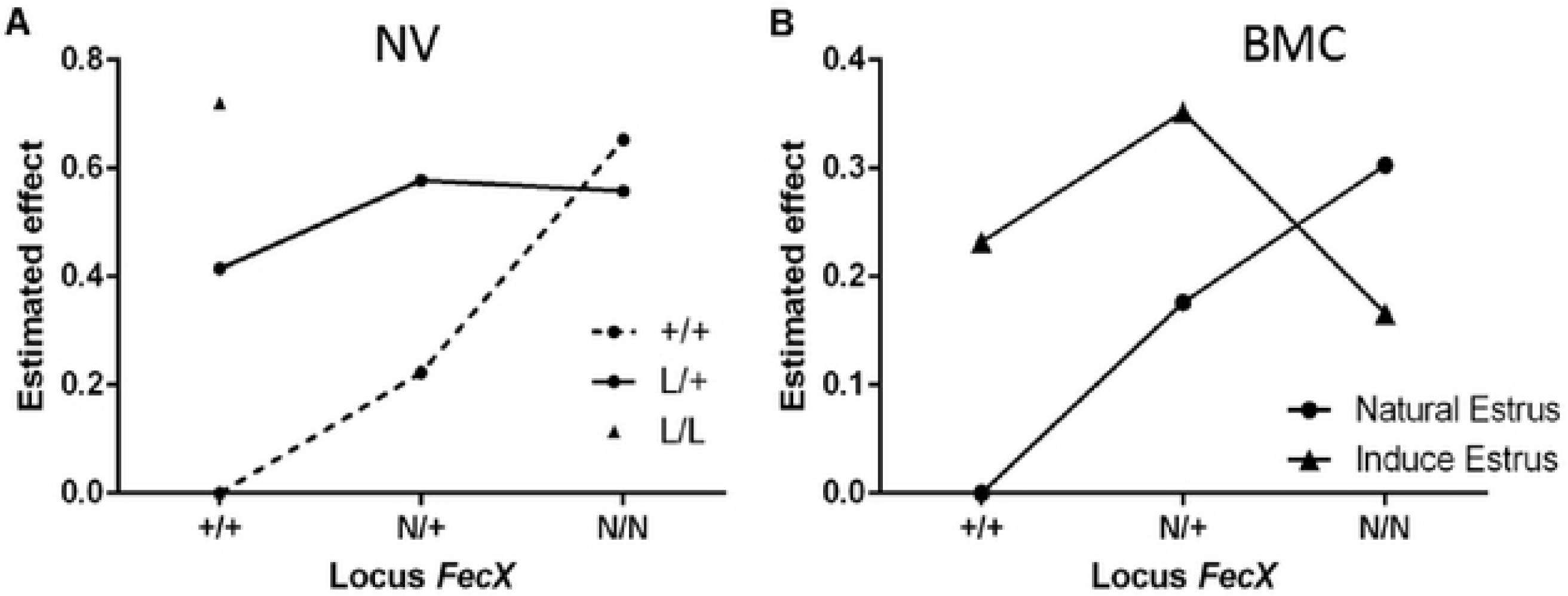
Estimated effect of *FecX*^*N*^. A) Estimated effect of *FecX*^*N*^ knowing *FecL*^*L*^ genotype in Noire du Velay (NV). B) Estimated effect of *FecX*^*N*^ according to induced or natural estrus in Blanche du Massif Central (BMC). Estimated increased LS from the linear mixed models with interaction between genotypes at the *FecX* and *FecL* loci for NV, and between *FecX* genotypes and estrus type for BMC are given relative to *L* and *N* alleles non-carrier ewes (+/+).

### Functional effects of the *FecX*^*N*^ mutation

As described above, *FecX*^*N*^ is located upstream of the coding region of the *BMP15* gene when referencing to the ovine genome v3.1 (ensembl.org) or v4.0 (ncbi.nlm.nih.gov). In both versions of the ovine genome, the *BMP15* gene annotation begins at the ATG start site and *FecX*^*N*^ is located −290pb upstream, possibly in the 5’UTR and/or the proximal promoter region. As a first approach, we took advantage of RNA sequencing data from ovine oocytes publicly available at EMBL-EBI (Bonnet et al., 2013[24]). After reads mapping against the ovine genome (v3.1) using STAR2 aligner within the Galaxy pipeline and visualization with Integrative Genome Viewer (IGV), the Fig 5 shows the location of *FecX*^*N*^ within the possible 5’UTR of the *BMP15* gene when expressed in the oocyte. Consequently, we have first tested the potential functional impact of *FecX*^*N*^ on the *in vitro* stability and translatability of the *BMP15* mRNA. Thus, the reference (T, *FecX*^+^) and variant (A, *FecX*^*N*^) forms of the ovine BMP15 cDNA (−297, +1183 referring to ATG start codon) were cloned in a pGEM-T vector for subsequent *in vitro* T7 promoter-dependent transcription/translation experiment using reticulocyte lysate solution. As shown in Fig 6, the western blotting of the BMP15 proteins produced from both forms and their chemiluminescent quantification revealed that the *FecX*^*N*^ mutation had no significant impact on the overall stability and translatability of the *BMP15* mRNA in this condition.

**Figure 5.**
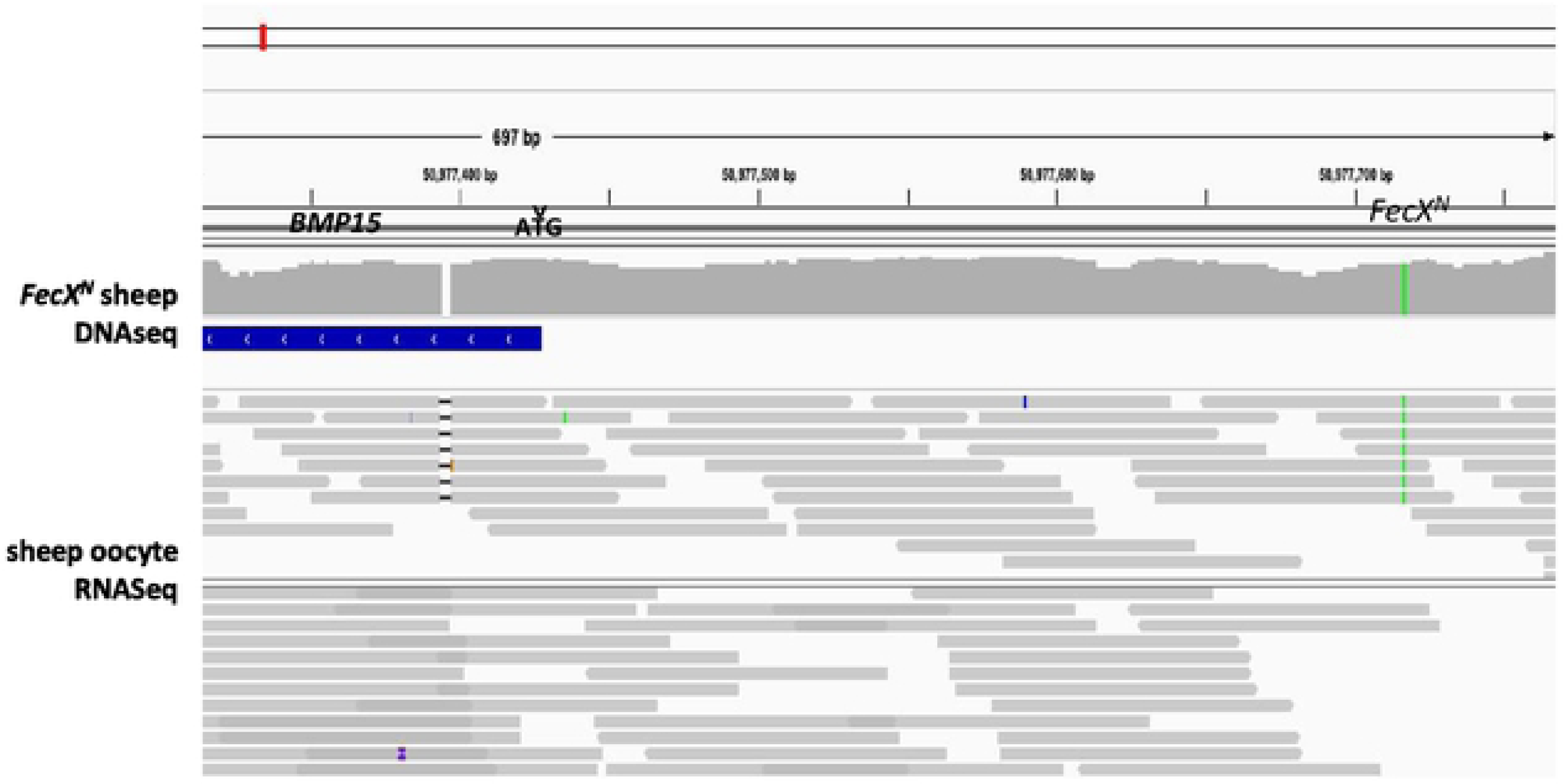
Localization of FecX^*N*^ in the *BMP15* upstream region. Integrative Genomics Viewer (IGV v2.4.10) snapshot of the ovine *BMP15* gene upstream region alignments (coordinates from OAR v3.1 assembly) with reads from whole genome DNA sequencing (DNASeq) of a homozygous *FecX*^*N*^ carrier ewe (green line) and total mRNA sequencing of ovine oocytes from small antral follicles (RNASeq, Bonnet et al. 2013), indicating a possible localization of *FecX*^*N*^ in the 5’UTR region of *BMP15* (ATG start codon is indicated by an arrowhead and *BMP15* gene annotation is on the minus strand).

**Figure 6.**
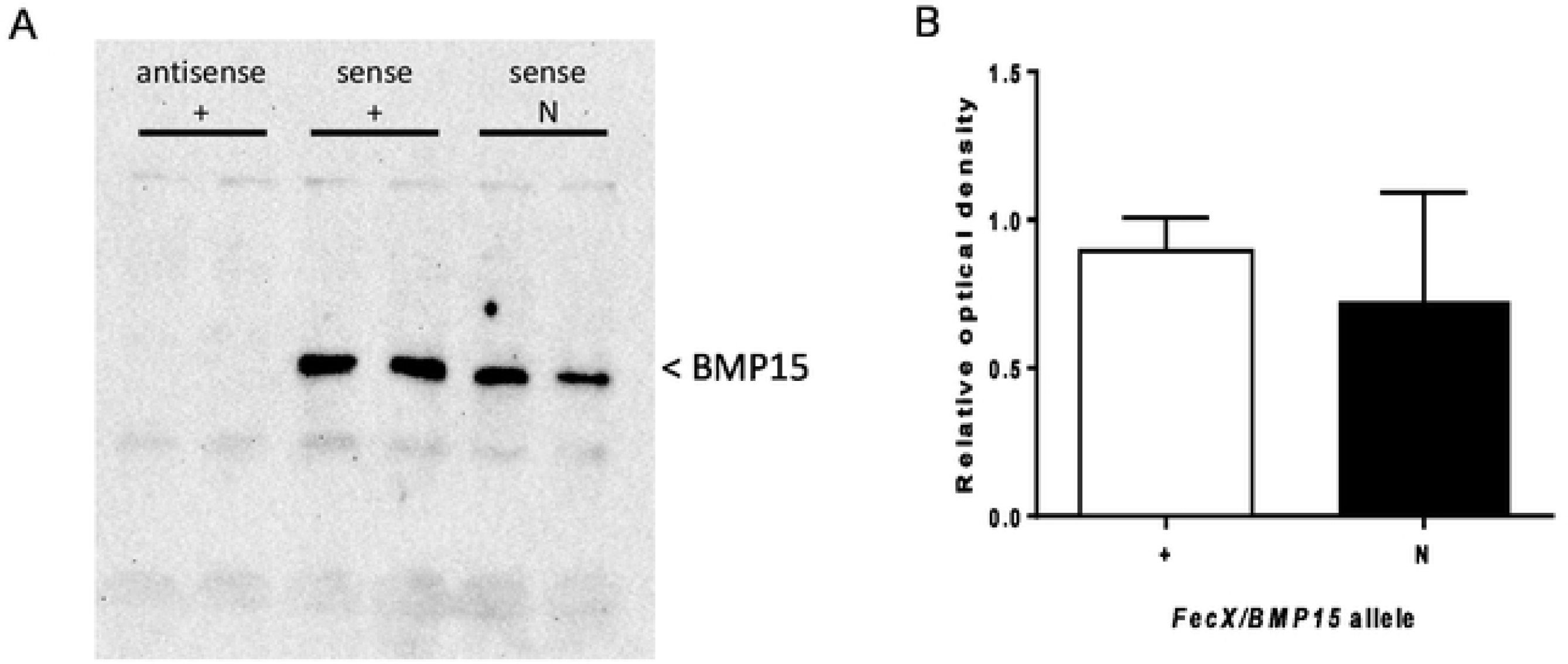
Effect of *FecX*^*N*^ mutation on the ovine BMP15 protein produced *in vitro.* *In vitro* transcription and translation were realized in duplicate in 6 independent experiments from antisense *BMP15* cDNA carrying the wild-type allele (antisense +) as negative control, or sense *BMP15* cDNA with (sense N) or without *FecX*^*N*^ (sense +). A) A representative western blot experiment revealing the BMP15 protein. B) Chemiluminescent signal of BMP15 was captured by a ChemiDoc MP imaging system and images were quantified (relative to sense +) and analyzed with the Image Lab Software (Bio-Rad).

As a second hypothesis, we have tested the *FecX*^*N*^ impact on the *BMP15* promoter activity. Two promoter regions were tested ([-743,-11] bp and [-443,-102] bp referring to ATG start codon) cloned in front of the luciferase reporter gene and transiently expressed in CHO cells cultured *in vitro*. As shown by the luciferase assays (Fig 7), the *FecX*^*N*^ variant was able to significantly reduce the luciferase activity in the context of both the long or the short BMP15 promoters, indicating the possible inhibitory impact of *FecX*^*N*^ on *BMP15* gene expression. To go further with this hypothesis, *in vivo* BMP15 gene expression was measured directly on isolated oocytes pools from NV and BMC homozygous ewe carriers and non-carriers of the *FecX*^*N*^ allele. Real-time qPCR experiments revealed a tendency of the *BMP15* expression to be decreased by 2-fold (*P*=0.17, genotype effect, two-way ANOVA) in the oocytes of *FecX*^*N*^ carriers despite a large inter-animal variability. In contrast, the expression of the second oocyte-specific prolificacy major gene *GDF9* seemed unaffected (Fig 8).

**Figure 7.**
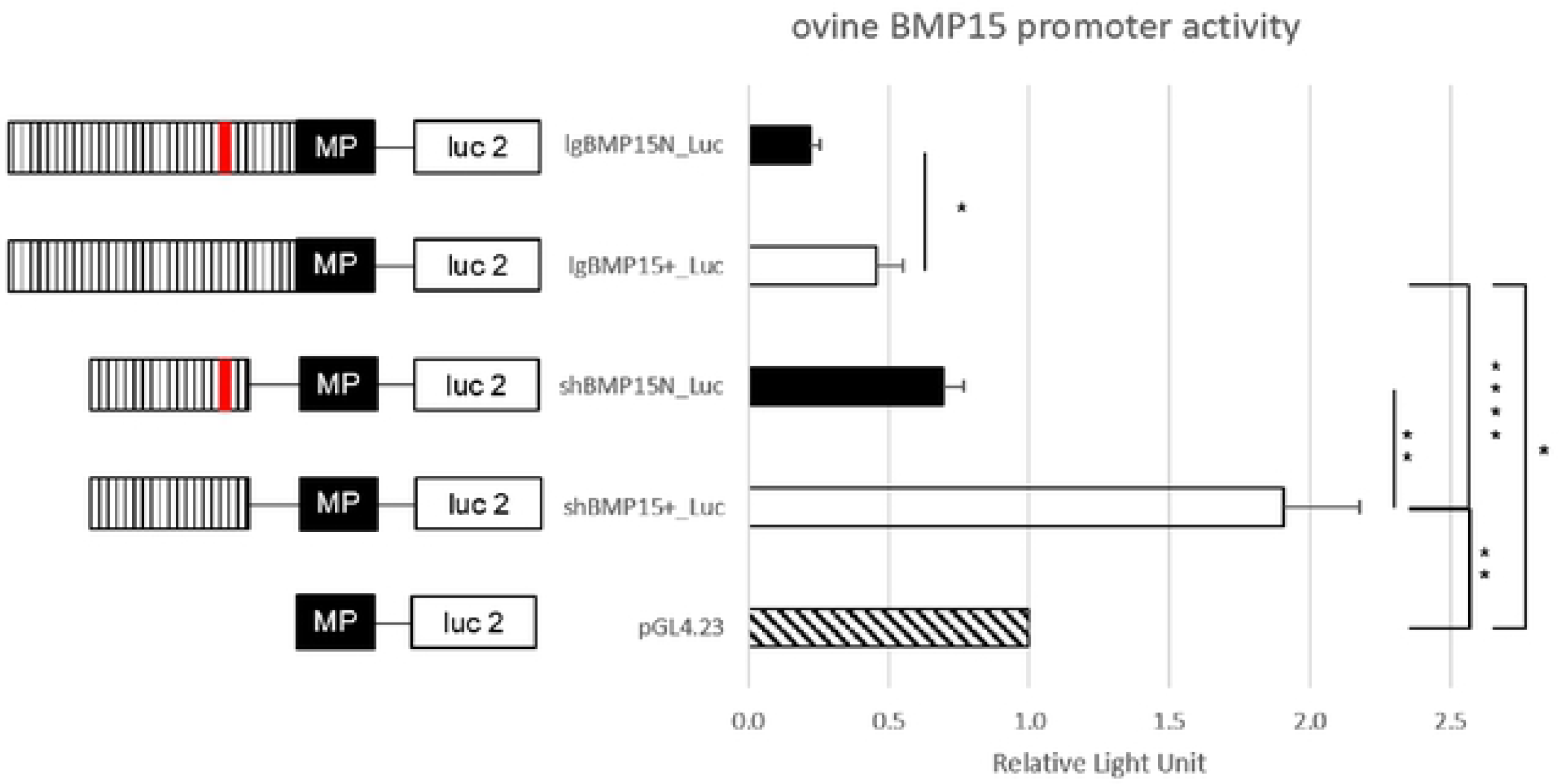
Functional effect of *FecX*^*N*^ mutation on the *BMP15* promoter activity. *In vitro* reporter luciferase assay from CHO cells transiently transfected with empty vector or wild-type *BMP15* promoter (BMP15^+^) or mutant *BMP15* promoter (BMP15^N^) Two fragments were generated, a long (lg) form of 732 bp (−743, −11bp referring to ATG start codon) and a short (sh) form of 341bp (−443, −102bp). Results are expressed as means ± SEM of the relative light unit (RLU) from 6 independent transfection experiments in triplicate. Asterisk indicates significant difference *: p<0.05; **: p<0.01; ****: p<0.0001.MP: minimal promotor of pGL4.23, Luc 2: luciferase reporter gene, hatched bar: *BMP15* promotor, red line: *FecX*^*N*^ mutation

**Figure 8.**
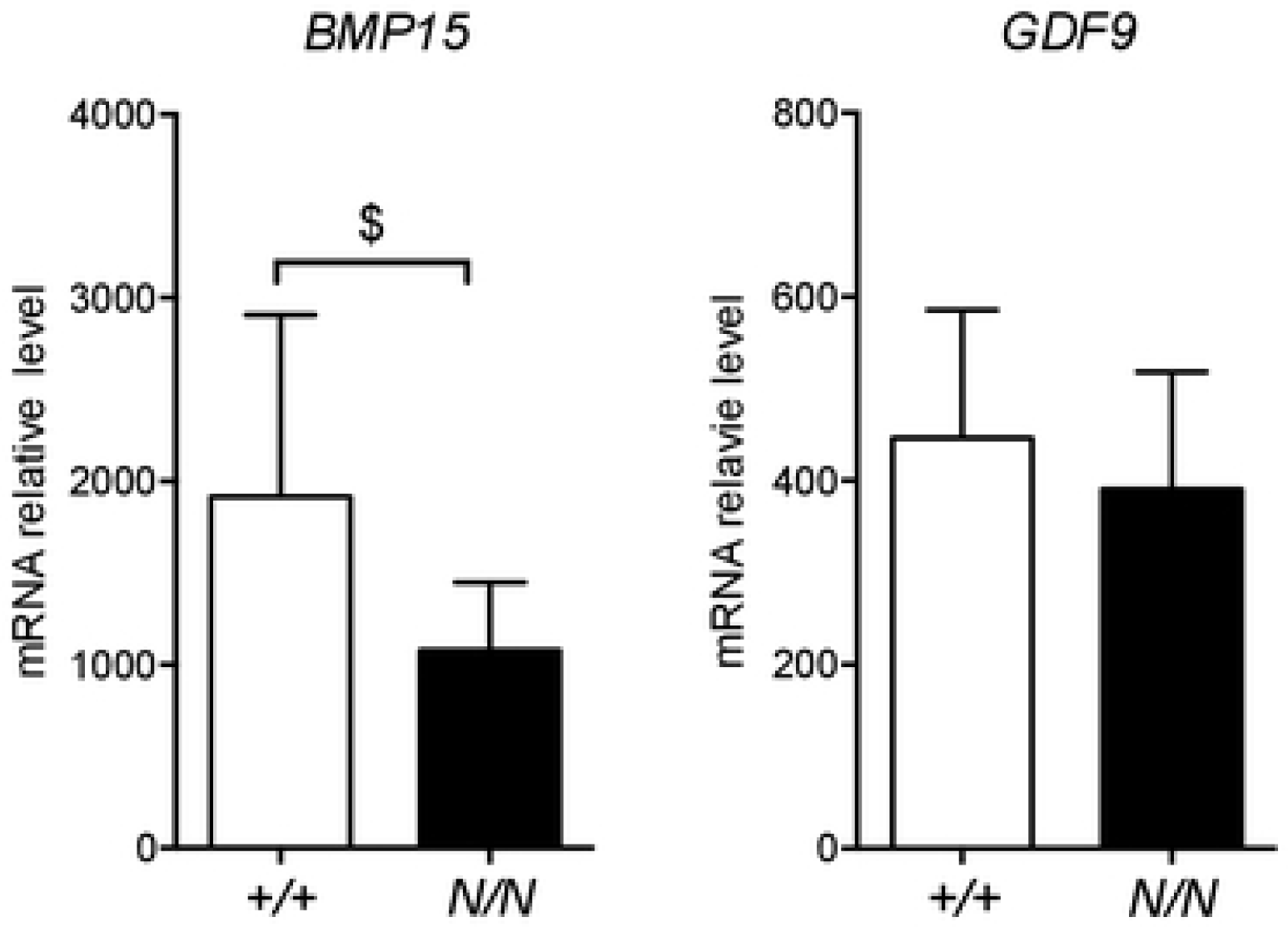
Effect of *FecX*^*N*^ mutation on *BMP15* and *GDF9* expression in ovine oocyte. Quantitative real-time PCR results of BMP15 (A) and GDF9 (B) expression in oocyte pools from growing (1-3mm) follicles (n=20 pools of 5 oocytes) from homozygous *FecX*^*N*^ carrier *(N*/*N*, n=S) and non-carrier ewes (+/+, n=S) during the follicular phase of the estrus cycle. Results are expressed as means ± SEM of the mRNA relative level for each genotype, using GAPDH and SDHA as internal references. Raw data were analyzed by two-way ANOVA. Dollar symbol indicates a suggestive difference between genotypes, $: p=0.166.

## Discussion

The present study identified the g.50977717T>A variant on the ovine chromosome X upstream of the *BMP15* gene as the most likely causative mutation for the increased prolificacy of the NV ewes. The highly significant genetic association with the extreme LS phenotype, the significant effect of the A variant on increasing prolificacy by +0.2 lamb per lambing in a large set of NV ewes, also found in the BMC genetic background, and the demonstrated action on *BMP15* transcriptional activity all support the causality of this mutation named *FecX*^*N*^.

The *BMP15* gene is at the top of the list of candidate genes controlling the ovarian function, ovulation rate and thus prolificacy in the ovine species, with nine independent causal mutations identified out of the sixteen already known. Indeed, 7 SNPs and 2 small INDELs all within the open reading frame were evidenced affecting the BMP15 function. Among these mutations, 2 SNPs and the 2 INDELS impaired the protein production either by generating premature stop codon (*FecX*^*H*^, Galloway et al. 2000; *FecX*^*G*^, Hanrahan et al. 2004 [11,12]) or by breaking the reading frame (*FecX*^*R*^, Martinez-Royo et al. 2008; *FecX*^*Bar*^, Lassoued et al. 2017[25,26]). The 5 other SNPs generate non-conservative amino acid substitutions all leading to a loss of function of BMP15 ranging from inhibited protein production (*FecX*^*L*^, Bodin et al. 2007 [17]), impaired interaction with GDF9 (*FecX*^*I*^ and *FecX*^*B*^, Liao et al. 2004 [27]), to altered cell signalling activity (*FecX*^*Gr*^ and FecX^O^, Demars et al. 2013 [14]). In contrast with the 9 mutations described above, the *FecX*^*N*^ variant evidenced in the present study is not located in the open reading frame of *BMP15* and does not alter the protein sequence. However, no other polymorphism genetically linked to *FecX*^*N*^ was found in the *BMP15* coding sequence when checked by whole genome or local Sanger sequencing of the *BMP15* gene from *FecX*^*N*^ carrier animals. Of course, this does not rule out the possibility of a polymorphism lying in another gene nearby with a still unknown role in the ovarian function and prolificacy. Nevertheless, we did not find any polymorphism (SNP and INDEL) altering the coding sequence of genes annotated in the significantly LS-associated genetic region of 3.5Mb on OARX (S1 Table), leaving *BMP15* as the most obvious candidate.

Whatever the version of the ovine reference genome (Oar_v3.1, Oar_v4.0 or even the last Oar_rambouillet_v1.0) the annotation of the *BMP15* gene always starts at the ATG initiating codon. Using publicly available transcriptome data from ovine oocytes RNAseq analysis, we were able to show that *FecX*^*N*^ located 290bp upstream of *BMP15* could stand in its 5’UTR region. From our *in vitro* functional analyses, *FecX*^*N*^ was not demonstrated to influence the translatability of the *BMP15* mRNA, but in the contrary it was shown to decrease the *BMP15* promoter activity. Little is known about transcription factors able to regulate *BMP15* expression. Several regulatory elements were evidenced in the pig *BMP15* promoter hosting consensus binding sites for LHX8, NOBOX and PITX1 transcription factors. However, only LHX8 was demonstrated as functionally activating the porcine BMP15 promoter activity (Wan et al. 2015 [28]). In human, a regulatory mutation in the 5’UTR of *BMP15* (c.-9C>G) was associated to non-syndromic premature ovarian failure (Dixit et al. 2006 [29]), but also to iatrogenic ovarian hyperstimulation syndrome (Moron et al. 2006 [6]). This mutation was shown to enhance the fixation of the PITX1 factor transactivating the BMP15 promoter (Fonseca et al. 2014 [30]). However, the *FecX*^*N*^ position does not fit with the syntenic location of porcine LHX8 and human PITX1 binding sites on the ovine *BMP15* promoter. Using the MatInspector promoter analysis tool (Genomatix), we were only able to hypothesize an alteration by *FecX*^*N*^ of a putative TATA-box like sequence (TTA<u>A</u>ATA >TTA<u>T</u>ATA). Unfortunately, our electromobility shift assay attempts using CHO nuclear extracts failed to demonstrate the binding of any factor at the *FecX*^*N*^ position, preventing us from defining the precise molecular mechanism by which *FecX*^*N*^ decreases the *BMP15* promoter activity.

The inhibition of the promoter activity combined with the apparent decreased of *BMP15* mRNA accumulation in homozygous *FecX*^*N*^*/FecX*^*N*^ oocytes seem to confirm the transcriptional regulatory role of *FecX*^*N*^. However, the moment we have chosen during the follicular phase of the late folliculogenesis for the comparative analysis between *FecX*^+^ and *FecX*^*N*^ oocytes from antral follicles could not be optimal to visualize a highly significant differential expression of *BMP15*. The *BMP15* gene expression in ovine oocytes begins during the primary stage of follicular development and its expression increases up to the antral stages (McNatty et al. 2005; Bonnet et al. 2011 [31,32]). Moreover, the streak ovaries phenotype of infertile ewes carrying homozygous mutations in *BMP15* have evidenced its crucial role in controlling the primary to secondary follicle transition (Galloway et al. 2000; Bodin et al. 2007; Lassoued et al. 2017 [11,17,26]). Consequently, it would certainly be appropriate to follow the *BMP15* expression in *FecX*^*N*^ carrier ewes from these early stages of folliculogenesis to better decipher the mutation impact on ovarian physiology. Nevertheless, the fact that *FecX*^*N*^ inhibits the *BMP15* gene expression fits well with the physiological and molecular models associating BMP system loss-of-function and increased sheep prolificacy (Fabre et al., 2006, Demars et al. 2013[14,33]).

One copy of *FecX*^*N*^ allele significantly increased by +0.30 to +0.50 the raw mean LS of NV ewes. When corrected for different environmental effects and more particularly for the genotype at the *FecL* locus, the estimated effect of *FecX*^*N*^ on LS was +0.22 lamb per lambing for the first copy and +0.43 for the second copy. This effect was in the range of already known prolific alleles in various sheep breeds (Jansson, 2014 [34]). The effect of *FecX*^*N*^ on LS seems independent of the genetic background. Indeed, the estimated positive effect of *FecX*^*N*^ on prolificacy was confirmed in BMC breed with +0.18 lamb per lambing based on natural estrus. Moreover, the same robust effect was observed even in the presence of PMSG for synchronizing the estrus cycles preceding the lambing. The same observation is made for other mutations controlling sheep prolificacy. For instance, the *FecL*^*L*^ allele exhibited a similar effect on LS in NV (+0.41, present study; +0.42, Chantepie et al. 2018 [15]), Lacaune (+0.47, Martin et al. 2014 [35]) and D’man (+0.30, Ben Jemaa et al. 2018 [22]), and this was also observed for the *FecB*^*B*^ allele introgressed in several populations (Kumar et al. 2008 [36]).

By genotyping a diversity panel, we also evidenced the presence of 2 *FecX*^*N*^ carrier animals in the Lacaune meat strain which will require further genotyping of numerous animals. If this is confirmed, the Lacaune meat breed will be another population, as Belclare, where 3 different natural prolific mutations are segregating (Hanrahan et al. 2004; Bodin et al. 2007; Drouilhet et al. 2013 [12,16,17]). The presence of *FecL*^*L*^ in both NV and Lacaune, and the presence of *FecX*^*N*^ in NV, BMC and Lacaune, also raises the question of the origin of these mutations. From population structure analysis, it was shown that NV, BMC and Lacaune shared the same origin within the European southern sheep populations that may explain the segregation of the same mutations in these populations (Rochus et al., 2018 [23]).

In conclusion, through a case/control GWAS strategy and genome sequencing, we have identified in the NV breed a second prolific mutation named *FecX*^*N*^ affecting the expression of the *BMP15* gene, a well-known candidate gene controlling OR and LS in sheep. This work confirms the relevance of the whole genome approaches to decipher the genetic determinism of the prolificacy trait. Homozygous *FecX*^*N*^/*FecX*^*N*^ animals were still hyperprolific as already observed for *FecX*^*Gr*^ and *FecX*^*O*^, but in contrast with sterile animals observed for the 7 other *FecX* homozygous variants in *BMP15*. As an upstream regulatory mutation, *FecX*^*N*^ also contrasts with these 9 other prolific causal mutations all evidenced in the coding part of *BMP15* and altering the protein function. Thanks to this new sheep model, the genetic etiology of ovarian pathologies in women could be improved by searching polymorphisms, not only in the coding region, but also in the regulatory parts driving the *BMP15* expression within the oocyte.

## Materials and Methods

### Animals

Ewes (*Ovis aries*) from the NV breed (n=2266) were genotyped on blood DNA at the *FecL* locus as already described (Chantepie et al. 2018[15]). In order to test the hypothesis of the segregation of a second major mutation controlling LS in this breed, a first set of 80 ewes with at least 5 LS records (mean LS=1.84; ranging from 1.00 to 3.50) were selected among the *FecL*^*+*^ homozygous genotype (n=2151, mean LS=1.58). Subsequently, for NV breed, the effect of the *FecX*^*N*^ mutation on LS was estimated on 2252 ewes, considering the genotype at the *FecL* locus. The presence of the *FecX*^*N*^ mutation in other breeds was checked on a diversity panel of 725 animals from 26 French sheep breeds (Rochus et al. 2018[23]; Table 3). For the BMC population, the effect of the *FecX*^*N*^ mutation on LS was estimated on 2456 ewes. For gene expression analysis, 10 homozygous ewes at the *FecX*^*N*^ locus (5 carriers and 5 non-carriers of the *N* allele) were bought from private breeders (6 NV and 4 BMC) and reared at INRA experimental facility (agreement number: D3142901). All experimental procedures were approved (approval number 01171.02) by the French Ministry of Teaching and Scientific Research and local ethical committee C2EA-115 (Science and Animal Health) in accordance with the European Union Directive 2010/63/EU on the protection of animals used for scientific purposes.

### Biological samples

All blood sampling from the numerous sheep breeds studied were collected from jugular vein (5 ml per animal) by Venoject system with EDTA and directly stored at −20°C for further use. Part of these blood samples (GWAS and diversity panel) was used for extraction of genomic DNA as described (Bodin et al. 2007[17]). All other samples were used for direct genotyping on whole blood without DNA purification (Chantepie et al. 2018[15]).

For ovary collection and oocyte isolation, the estrus cycles of all adult NV and BMC ewes were synchronized with intravaginal sponges impregnated with flugestone acetate (FGA, 30 mg, CEVA) for 14 days. Ovaries were collected at slaughtering during the follicular phase 36h after FGA sponge removal. Cumulus-oocyte complexes (COC) were immediately recovered from all visible 1-3mm follicles by aspiration using a 1ml syringe with a 26G needle and placed in McCoy’s 5A culture medium (Sigma-Aldrich). COC were mechanically dissociated by several pipetting and washing cycles in 150µl drops of McCoy’s 5A medium and finally, denuded oocytes devoid of granulosa cells were recovered in 1X PBS. Only intact oocytes with a good homogeneity of the cytoplasm were grouped to obtain two to three pools of 5 oocytes per animal and stored at −80°C before RNA extraction.

### Genotyping analyses

The *FecL*^*L*^ mutation (OAR11:36938224T>A, NC_019468) was genotyped directly on whole blood samples by the KAPA-KASP assay as already described (Chantepie et al. 2018[15]). As a prerequisite before GWAS, a set of 30 high prolific *FecL*^*+*^*/FecL*^*+*^ ewes were controlled for the absence of other evidenced major mutations affecting sheep prolificacy in French populations. Using the same KAPA-KASP assay, *FecX*^*L*^ and *FecX*^*Gr*^ alleles in *BMP15* were genotyped as described (Chantepie et al. 2018[15]). *FecB*^*B*^ in the exon 7 of the *BMPR1B* gene (OAR6:29382188A>G, NC_019463.1) was genotyped using forced restriction fragment length polymorphism (RFLP) as described by Wilson et al. (2001)[18].

The whole genome genotyping was performed on 80 ovine genomic DNA using the OvineSNP50 Genotyping Beadchip from Illumina according to the manufacturer’s protocol at the Laboratoire d’Analyses Génétiques pour les Espèces Animales, (LABOGENA, Jouy en Josas, France; www.labogena.fr). From the dataset, individuals with a call rate <0.98 were excluded. SNP exclusion thresholds were: call frequency <0.95, and minor allele frequency (MAF) <0.01; or a significant deviation from Hardy-Weinberg equilibrium (HWE) in the controls (p<1.10^−6^). Non-polymorphic SNP positions and markers with no position on the OARv3.1 reference genome map were also discarded. Finally, from the available design of 54241 SNPs available on the Illumina OvineSNP50 Beadchip and 80 selected NV ewes, the final dataset was reduced to 47446 SNPs analyzed in 79 individuals.

The *FecX*^*N*^ mutation (OARX: 50977717T>A, NC_019484) was genotyped by a RFLP analysis using the Mse1 restriction enzyme (New England Biolabs) after a first step of Terra PCR Direct Polymerase Mix amplification (Takara) using one μl sample of total blood. The accuracy of the *FecX*^*N*^ RFLP genotyping was controlled by Sanger sequencing on few samples using the same amplification primers.

PCR amplifications were conducted independently for each locus studied on an ABI2400 thermocycler (Applied Biosystems) with the following conditions: 5min at 94°C, 32 cycles of 30s at the specific melting temperature, 30s at 72 °C and 30s at 94 °C, followed by 5 min at 72 °C. The primers used in this study are listed in S2 Table.

### Whole genome sequencing (WGS) analysis

DNA sequencing libraries were constructed from 1 µg of genomic DNA using TruSeq DNA PCR-free Library Prep kit (Illumina). Sequencing was run on an Illumina HiSeq 2500 apparatus using a paired-end read length of 2×150 pb with the Illumina Reagent Kits as already described (Demars et al. 2017[37]). WGS was performed at the Genotoul-GeT core facility (INRA Toulouse, https://get.genotoul.fr). The raw reads of Illumina DNA sequencing were preprocessed by removing adapter sequences. After quality control, the FastQ files and metadata were submitted to the European Nucleotide Archive (ENA) at EMBL-EBI (accession number PRJEB35553). Reads mapping and variants calling were performed using the local instance of Galaxy (https://galaxyproject.org) at the Toulouse Midi-Pyrénées bioinformatics platform (http://sigenae-workbench.toulouse.inra.fr). The cleaned paired reads were combined and mapped against the ovine genome assembly (Oar_v3.1.86) using BWA-MEM (Galaxy version 0.7.17.1). The resulting BAM files were sorted using Samtools_sort (Galaxy version 1.0.0). Sorted and indexed BAM files were visualized through Integrative Genome Viewer, IGV software version 2.4.10 (Robinson et al. 2011[38]). A GFF3 annotation file was obtained from Ensembl (Ovis_aries.Oar_V3.1.78). We applied GATK version 3.5-0 to performed SNP and InDel discovery and genotyping across the two samples simultaneously using standard filtering parameters according to GATK Best Practices recommendations (DePristo et al. 2011; Van der Auwera et al. 2013[39,40]). Variants effect and annotation were realized by SNPEff version 4.1 and filtering of interesting variants was performed using the SNPSift tool.

### In vitro transcription and translation of BMP15

The full-length cDNA of ovine BMP15 (1480bp [-297,+1183] referring to ATG start codon) with or without the *FecX*^*N*^ mutation was generated from oocyte-derived RNA after a reverse transcription (RT) step (described in RNA extraction and RT paragraph, primers are listed in S2 Table). The resulting PCR products were inserted by TA cloning into pGEM-T Easy plasmid (Promega) possessing T7 and SP6 promoters. The orientation of insertion and exclusion of unexpected PCR-induced mutations were controlled by Sanger sequencing.

*In vitro* transcription and translation were realized from 500ng of cDNA pGEM-T construct using the TnT T7 Quick Coupled Transcription/Translation kit (Promega) and Transcend Biotin-Lysine-tRNA following the manufacturer protocol. Reactions for each construct were run in duplicate in 6 independent TnT experiments. The resulting BMP15 protein was revealed using Transcend non-radioactive translation detection system with chemiluminescent method (Promega) after reducing SDS-PAGE on a gradient (4-15%) polyacrylamide gel (Promega) and transfer onto nitrocellulose membrane. Chemiluminescent signal was capture by a ChemiDoc MP imaging system and images were analyzed with the Image Lab Software (Bio-Rad).

### BMP15 promoter activity

The promoter sequence of the ovine *BMP15* gene was amplified by PCR on genomic DNA from both homozygous *FecX*^+^ and homozygous *FecX*^*N*^ ewes. Two sizes of fragments were generated for cloning the *BMP15* promoter in front of the luciferase (Luc) reporter gene, a long (lg) form of 732 bp ([-743,-11] bp referring to ATG start codon) and a short (sh) form of 341bp ([-443,-102] bp). The PCR products were engineered for digestion using Kpn1 and Hind3 restriction enzymes (New England Biolabs) and inserted into the pGL4.23 vector (Promega). The four resulting constructs (lgBMP15^+^-Luc; lgBMP15^N^-Luc; shBMP15^+^-Luc; shBMP15^N^-Luc) were controlled by Sanger sequencing. Primers used to generate these constructs are listed in S2 Table. Twenty-four hours after seeding (3.10^4^ cells/well, 24 wells plate), CHO (Chinese Hamster Ovary) cells were transfected using Lipofectamine 3000 (Invitrogen) with 500ng/well of pGL4.23 constructs either empty or containing *BMP15* promoter fragment. Forty-eight hours after transfection, cells were lysed and assayed for luciferase activity (Luciferase reporter assay kit, Promega). Luminescence in relative light units (RLU) was measured by a Glomax microplate reader (Promega). Each construct was assayed in triplicate in 6 independent transfection experiments.

### RNA extraction, reverse transcription and quantitative PCR

Total RNA from pools of 5 oocytes were extracted using the Nucleospin RNA XS kit according to the manufacturer’s protocol (Macherey-Nagel) and including a DNase1 treatment. The low quantity of RNA recovered did not allow quantification. So, the equivalent of 1.25 oocyte was reverse-transcribed using SuperScript II reverse transcriptase (Invitrogen) and anchored oligo(dT)22 primer (1µl at 10 µM). Primer design using Beacon designer 8.20 (Premier Biosoft), SYBR green real-time PCR cycling conditions using QuantStudio 6 Flex Real-Time PCR system (ThermoFisher Scientific) and amplification efficiency calculation (E=e^(−1/slope)^) were as already described in Talebi et al. (2018)[19]. Primer sequences, amplicon length and amplification efficiency are listed in S2 Table. RNA transcript abundance was quantified using the ΔCt method with the mean expression of GAPDH and SDHA as internal references and following the formula R=[E_ref_^Ct ref^/E_target_ ^Ct target^]. The two reference genes were validated by the Bestkeeper algorithm (Pfaffl et al., 2004)[41].

### Data analysis

Single-marker association analyses were conducted using a Fisher’s exact test and a Bonferroni correction has been applied to check for significance levels. The chromosome-wide and genome-wide values have been established as mentioned by Balding et al. 2006 [42]. Statistical analyses were done using PLINK1.9 software under a case/control design [43]. Among the 79 datasets of 47446 SNPs analyzed, the LS trait was considered as case when mean LS ≥ 2.18 (n=39) and control when LS ≤ 1.45 (n=40). Haplotypic association analysis on X chromosome were performed using FastPhase software [44]. Empirical significance levels were calculated using maximum statistic permutation approach (max (T), n=1000).

Allele effect on LS was estimated in NV and BMC breeds on data extracted from the French national database for genetic evaluation and research managed by the Institut de l’Elevage (French Livestock Production Institute) and the CTIG (Centre de Traitement de l’Information Génétique, Jouy-en-Josas, France). Only females born after 2000 were retained (27 754 NV ewes with 122 110 LS records and 110 848 BMC ewes with 461 405 LS records) with their pedigree over 5 generations. *FecX*^*N*^ genotype effect on the subset of 79 case/control animals was assessed by one-way ANOVA, follow by Newman-Keuls post-hoc test. For the large animal cohort analyses, the linear mixed models used were as similar as possible to those of the national genetic evaluation system (Poivey et al. 1995 [45]). In the present study, the following fixed effects were considered: i) the genotype at the *FecX* locus, ii) the month of birth (12 levels) iii) a physiological status effect combining parity, age at first lambing, rearing mode and postpartum interval (44 levels) and iv) a combination of the flock year and season effect. Two random effects were added to the model: a permanent environmental effect and an animal additive genetic effect. Moreover, an additional fixed effect of the reproduction type was considered for the BMC breed for which some hormonal treatments are used each year (87% and 13% after natural and induced estrus in the data set). For the NV breed, since the *FecL*^*L*^ allele is also segregating in the population (Chantepie et al. 2018 [15]), the effect of the genotype at this locus (2252 known and 25502 unknown genotypes) as well as its interaction with the genotypes at the *FecX* locus were considered. All these models were fitted using the ASReml software (Gilmour et al. 2009 [46]).

The comparison between *FecX* alleles for BMP15 protein quantification was analyzed using Student’s t-test, using Welch’s correction. For reporter luciferase assays, differences between constructs were analyzed by one-way ANOVA followed by Newman-Keuls post-hoc test. QPCR data for *BMP15* and *GDF9* expression in oocytes were analyzed by two-way ANOVA considering genotype and breed effects. P > 0.05 was considered as not significant. All these experimental data are presented as means ± SEM and were analyzed using Prism 6 (GraphPad Software Inc.).

## Acknowledgments

We thank Claire Chantaduc, Didier Cathalan and Kévin Chile from ROM Sélection managing the NV and BMC populations, for their precious help in the planning of blood sampling. We are grateful to the breeders who made their animals available for this study. LC was supported by a PhD grant co-funded by APIS-GENE through the Proligen project and the European Funds for Regional Development (FEDER) through the Interreg POCTEFA programme in the framework of the PIRINNOVI project (EFA103/15). Part of the NV sampling was supported by the DEGERAM project co-funded by the FEDER Massif Central, the Régions: Aquitaine, Midi-Pyrénées, Limousin and Auvergne; and the French government.

## Supporting information captions

S1 Table. List of variants found in the OARX: 50639087-54114793 region. Listing of 60 SNPs and 90 small INDELs with quality score >30.

S2 Table. List of primers used in the study. Locations of primers are based on the OARv3.1 ovine genome assembly available on ensembl.org.

S3 Figure. Genome-wide and chromosome-wide association results integrating the SNP OARX: 50977717T>A. (A) Genome-wide association results for litter size in the NV sheep population. Manhattan plot shows the combined association signals (-log_10_(p-value)) on the y-axis versus SNPs position in the sheep genome on the x-axis and ordered by chromosome number (assembly OARv3.1). Red line represents the 5% genome-wide threshold. (B) OARX chromosome-wide association results. The curve shows the combined association signals (-log_10_(p-value)) on the y-axis versus SNPs position on the X chromosome on the x-axis (assembly OARv3.1). Red line represents the 5% chromosome-wide threshold. In both panels, the position of the SNP OARX:50977717T>A is indicated by a red dot. In panel (B), the *BMP15* gene location is indicated by a red arrowhead.

